# Domestication reprogrammed the budding yeast life cycle

**DOI:** 10.1101/2020.02.08.939314

**Authors:** Matteo De Chiara, Benjamin Barré, Karl Persson, Amadi Onyetuga Chioma, Agurtzane Irizar, Joseph Schacherer, Jonas Warringer, Gianni Liti

**Affiliations:** Université Côte d’Azur, CNRS, INSERM, IRCAN, Nice, France; Department of Chemistry and Molecular Biology, Gothenburg University, Gothenburg, Sweden; Department of Microbiology, University of Nigeria, Nsukka, Nigeria; Université de Strasbourg, CNRS, GMGM UMR 7156, Strasbourg, France

**Author notes:** These authors contributed equally to this work (Matteo De Chiara, Benjamin Barré). These authors jointly supervised this work (Jonas Warringer, Gianni Liti).

## Abstract

Domestication of plants and animals is the foundation for feeding the world population. We report that domestication of the model yeast *S. cerevisiae* reprogrammed its life cycle entirely. We tracked growth, gamete formation and cell survival across many environments for nearly 1000 genome sequenced isolates and found a remarkable dichotomy between domesticated and wild yeasts. Wild yeasts near uniformly trigger meiosis and sporulate when encountering nutrient depletions, whereas domestication relaxed selection on sexual reproduction and favoured survival as quiescent cells. Domestication also systematically enhanced fermentative over respiratory traits while decreasing stress tolerance. We show that this yeast domestication syndrome was driven by aneuploidies and gene function losses that emerged independently in multiple domesticated lineages during the specie’s recent evolutionary history. We found domestication to be the most dramatic event in budding yeast evolution, raising questions on how much domestication has distorted our understanding of this key model species.

## Introduction

Humans have domesticated plant and animal species over at least 12000 years to improve agricultural traits. Domestication greatly contributed to human population expansion and development by releasing labour from food production, but also profoundly altered the domesticated species. Controlled breeding and artificial selection on the segregating genetic diversity created plant and animal crops that now differ drastically from those of their ancestors and wild relatives^1^. Selected traits recur across domesticated species^2^ and, due to pleiotropy and linkage, often come with specific, non-desired side-effects. Human-created niches may also impose quasi-domestication and unintentionally favour some traits and genetic variants with side-effects linked to them^3^. Both domestication and quasi-domestication therefore results in suites of co-evolving traits, known as domestication syndromes^4,5^.

Domestication of the budding yeast, *Saccharomyces cerevisiae*, dates back over 9000 years, with the earliest use in rice, honey and fruit fermentation^6^. Subsequent domestication for industrial or semi-industrial beer, dairy, rice, cocoa, coffee^7^, bread, palm sap and agave fermentation^8^ transformed wild yeast lineages into specialized domesticated breeds^9–15^. Lab domestication and the subsequent use of budding yeast as a model organism resulted from efforts to breed pure, industrial beer strains^16^. Modern lab strains are genetic mosaics with genomes composed of DNA from diverse domesticated origins^17^. The continued existence of wild yeast not spoiled by extensive gene flow from feral domesticated yeast became accepted only recently^18,19^. Wild strains are more diverse in terms of single nucleotide variation; domesticated lineages are instead distinguished by genome content variation in form of aneuploidies, polyploidies and the copy number of individual genes and chromosome segments^20–23^. These structural and copy number variants in domesticated lineages may explain the high phenotypic variation among domesticated lineages and the unexpectedly large intraspecies phenotypic diversity in the species^9,20,21,24^.

While the impact of domestication on the yeast genome is well described, its effect on the yeast biology is poorly understood. The life cycle of an organism represents the very core of its organismal properties, directly defining its fitness. Life cycle changes are therefore dramatic evolutionary events with far-reaching consequences. Budding yeast reproduces asexually under nutrient abundance, expanding its population size. Growing cells first privatize and use the most easily catabolized carbon and nitrogen by repressing mitochondrial respiration and the uptake and use of non-preferred sources^25^. When the preferred sources are consumed, the repression is relaxed and yeast switches to non-preferred sources, including by respiring the produced ethanol. When an essential nutrient becomes scarce, cell growth and the asexual population expansion ceases. Diploid cells then either pass through meiosis and enter a haploid spore state^26^, shift from unicellular to multicellular filamentous organization^27^ or enter into a quiescent G0 stage^28^. These transitions favour survival by repressing metabolism, reproduction and translation, mobilizing nutrients intracellularly by autophagocytosis, and protecting against external challenges^29–33^. When the nutrient status switches back to favourable, cells re-activate translation, rebuild growth-required cell functions, revert to single, vegetative cell status and re-initiate the asexual cell cycle^34^.

We tracked asexual reproduction, entry into spore state and survival as quiescent cells across many environments for nearly 1000 genome sequenced *S. cerevisiae* strains^20^ and found a remarkable dichotomy between domesticated and wild yeasts. We found that domestication shifted yeast from respiratory to fermentative asexual reproduction and decreased resistance to stresses other than those associated with domestication environments. Concomitantly, domestication abolished or severely impaired the sexual life cycle and the capacity to enter into the protective haploid spore state. We traced genetic change driving this extraordinary yeast domestication syndrome to lineage-specific combinations of aneuploidies and loss-of-function mutations.

## Results

### Domestication promoted fermentative over respiratory growth and lowered stress resistance

We first revisited published growth yields for 971 wild and domesticated budding yeast strains that reproduced asexually in 36 stress environments^20^. Wild yeasts better tolerated most stresses (*n*=16, **Supplementary fig. 1a**; **Supplementary table S1**, FDR (False Discovery Rate), *α*<0.05), including heat, oxidative and osmotic stresses, while growing less well in those (*n*=5, **Supplementary fig. 1b**) that dominate in known domestication niches, such as sodium^3^, copper^35^ or arsenic^36^ exposure.

We next measured doubling times and yields of these yeasts while using common carbon and nitrogen sources^19,37^ (**Fig. 1, Supplementary data S1**). Domesticated yeasts were better adapted to maltose, sucrose and higher concentrations of glucose (>0.5%), fructose (>0.5%) and xylose (>2%); all carbon sources that characterize human-made fermenting niches^38^. Domesticated yeasts also better used isoleucine and valine, as well as threonine, which can be catabolized into isoleucine. Isoleucine and valine are degraded in the fermentation induced^39^ Ehrlich pathway by Bat1 and Bat2, while threonine is also anaerobically catabolized to propionate during fermentation. Wild yeasts were better adapted to most respiratory carbon sources (ethanol, glycerol) as well as to lower concentrations (up to 1%) of sugars; lower sugar concentrations promote respiration by relieving the glucose repression of respiratory genes^40^. Wild yeasts were also better at using citrulline, serine and alanine. The metabolism of these compounds produces fumarate or pyruvate (and through the latter: oxaloacetate), which are Krebs cycle intermediates intrinsically linked to respiration. In addition, since the replenishment of a cyclic pathway can be achieved by any anaplerotic reactions, the net result of the production of any of the Krebs cycle intermediates would entail a larger availability of precursors for other paths (such as the gluconeogenesis or the mitochondrial production of ATP)^41^. Wild yeasts also better catabolized glycine and phenylalanine.

**Figure 1.**
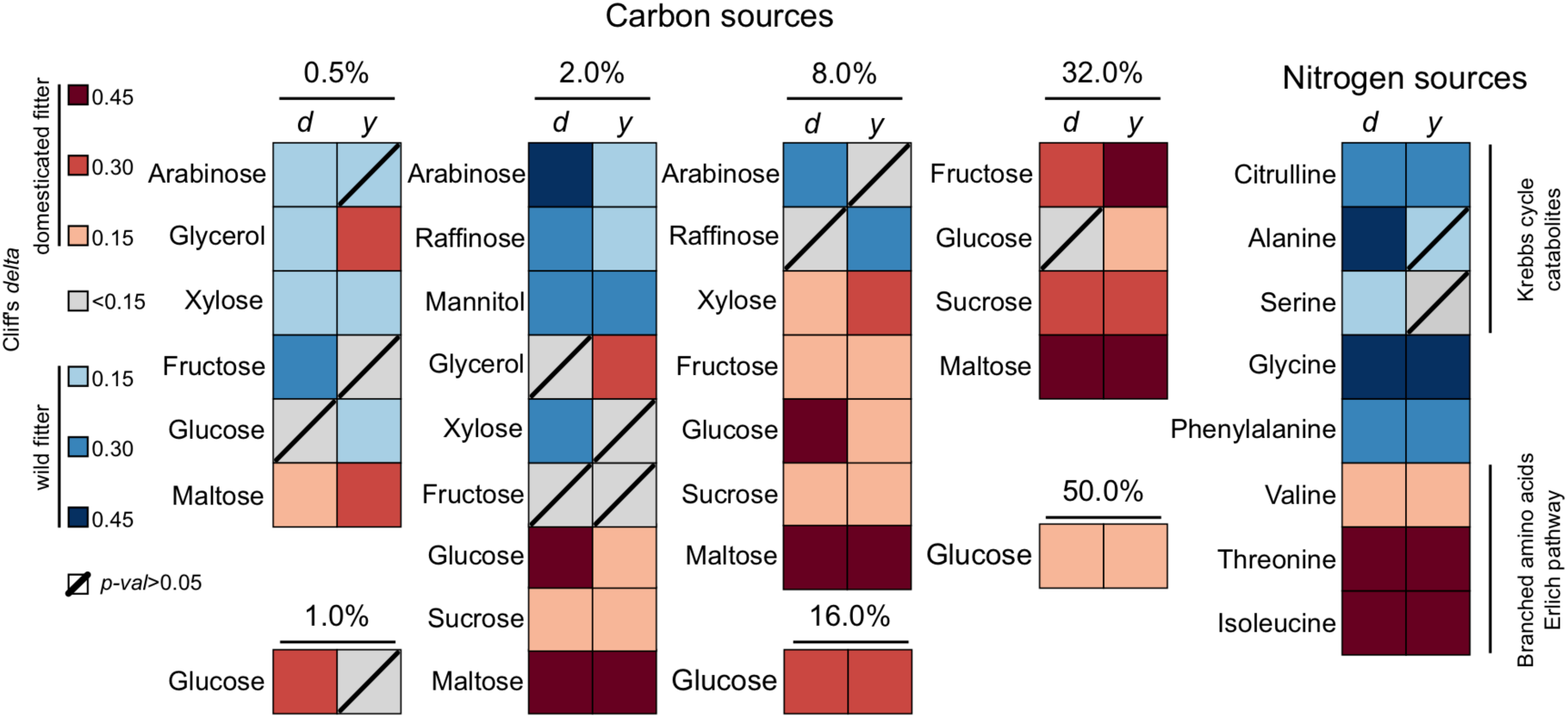
Domestication promoted fermentative over respiratory growth. Significant differences (Mann-Whitney: FDR *α*=0.05) in asexual growth between wild and domesticated yeast strains. Growth was measured as the log_2_ doubling time (*d*) and yield (*y*) of asexually reproducing, isogenic cultures. Colour intensity indicates increasing difference. Red = domesticated strains better. Blue = wild strains better. Crossed square: not significant.

We finally traced carbon and nitrogen adaptations down to 853 variants, by GWAS (**Supplementary data S1**). The vast majority (96%, *n*=815) of these was private to either doubling time (*d*) or yield (*y*), consistent with the very low correlation (*r*^2^ mean=0.16) of these fitness components across strains. We therefore conclude that the efficiency and rate of nutrient use likely evolved independently in budding yeast, and are independently regulated. We recovered known causes of domestication effects, e.g. *SUC2* gain (sucrose use, *d*), *MAL* gene amplification (maltose use, *d*), *DAL5* gain (citrulline use, *d*). We also disclosed many unknown effects, many of which were highly pleiotropic. Thus, yields in sugar-restricted niches often associated to the mitochondrial DNA polymerase *MIP1* or the cytochrome C peroxidase *CCP1*, whereas the corresponding doubling times often associated to either *SEC11* or *COG2* dependent intracellular transport. We conclude that variation in yeast use of carbon and nitrogen are in line with domestication selecting for fermentative over respiratory growth.

### Domestication abolished or impaired yeast sporulation

Yeast sporulation is known to be most efficiently induced by nitrogen starvation combined with respiratory growth^42,43^. We found wild yeasts (*n*=54) to sporulate rapidly in these conditions, while domesticated yeasts (*n*=503) were much slower (**Fig. 2a**). In particular, the domesticated French Dairy, African beer and Sake clades sporulated very poorly, while the Wine/European *S. boulardii* subclade completely failed to sporulate (**Supplementary fig. 2a and b**). Yeast in clades of unassigned type (*n*=293) sporulated as domesticated yeasts. Among unassigned yeasts, French Guiana strains that colonize human bodies were virtually unable to sporulate (**Supplementary fig. 2a**). We also determined how long the produced spores survives, by monitoring the spore lifespan of a wild (North American - AKN) and a well-sporulating domesticated (Wine/European - BTN) yeast. We found spore survival to remain amazingly high (>90%) without any detectable gamete mortality during the entire 8 weeks test period (**Fig. 2b**).

**Figure 2.**
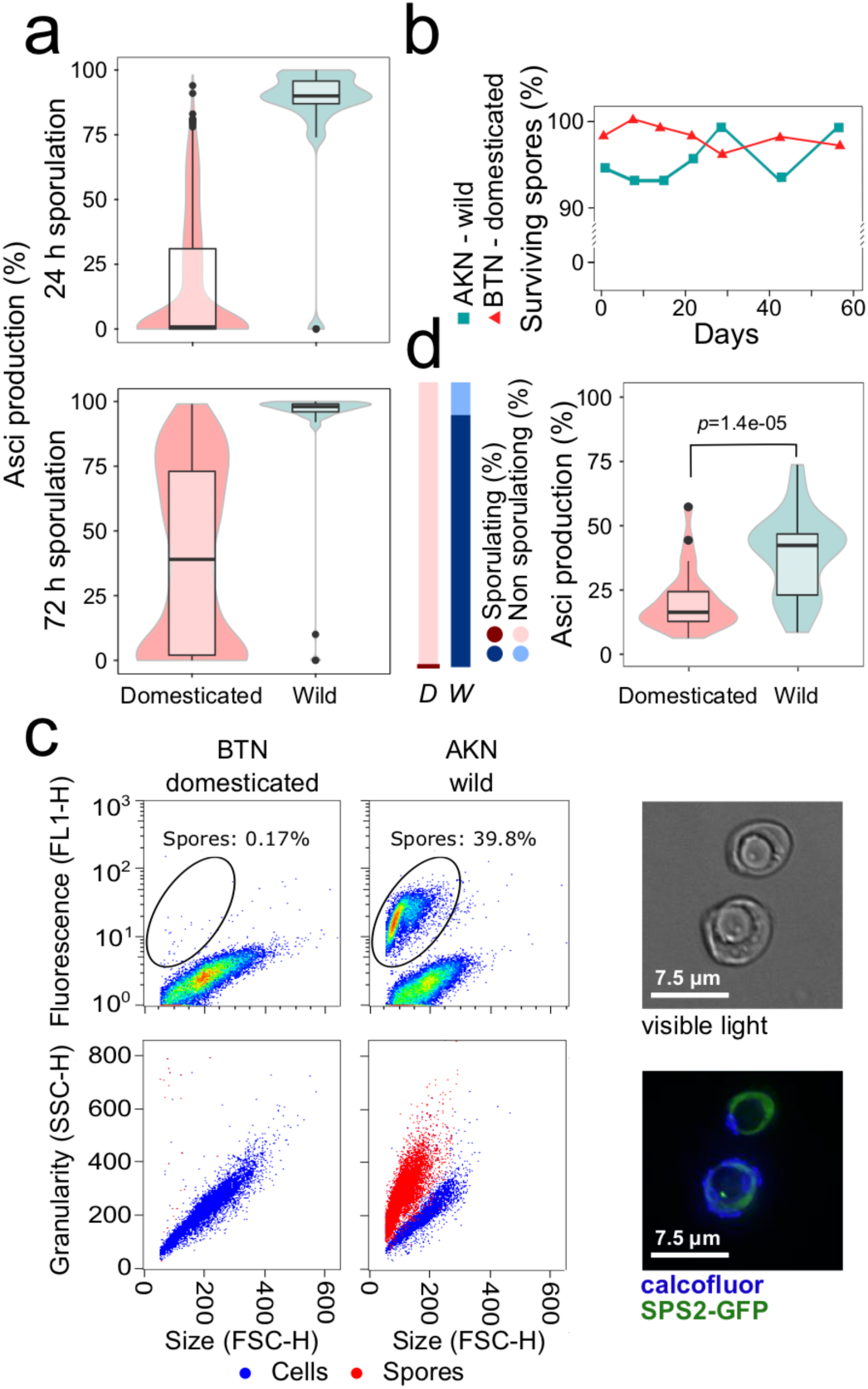
Domestication abolished or impaired yeast sporulation. **a**, Sporulation efficiency of wild and domesticated yeast in the traditional experimental sporulation (KAc) environment, as measured by microscopic counting of asci after 24 h (*upper panel*) and 72 h (*lower panel*). Box: interquartile range (IQR). Whiskers: 1.5x IQR. Thick horizontal line: median. **b**, Spore viability in water as a function of time, in one domesticated (BTN) and one wild (AKN) sporulating isolate. The fraction of viable spores was estimated as spores capable of forming visible colonies after 3 days on YPD medium. **c**, Validating a novel, FACS based analysis of sporulation in water, where asci only contain single (monads) or double (dyads) spores. A *SPS2*-GFP reporter construct was inserted in the monad producing wild strain AKN (*right panels*), which was used as positive control. We found p*SPS2* to be activated in spores only. A non-sporulating domesticated strain BTN (*left panels*), with the same *SPS2*-GFP reporter, was used as negative control. Sporulation was estimated after 8 days in water. *Upper panels:* Green fluorescence (FL1-H) vs. cell size (FSC). *Lower panel:* Cell granularity (SSC) vs. cell size (FSC), red = fluorescent cells, blue = non-fluorescent cells. *Right:* Normal (*upper*) and fluorescence (*lower*) microscopy of monads. **d**, FACS based analysis of the sporulation of domesticated and wild yeast after 8 days in water. *Left panel:* Fraction of domesticated (*D, n*=486) and wild (*W, n*=52) isolates sporulating. Fisher’s exact test: *p*=4.9e-39. *Right panel:* Sporulation efficiency in sporulating domesticated (*n*=26) and wild (*n*=46) isolates. Mann-Whitney: *p*=1.4e-05. Box: IQR. Whiskers: 1.5 X IQR.

We next investigated the ability of wild yeasts to sporulate under the most extreme starvation, i.e. in water. We developed a high-throughput flow cytometry assay based on cell size and granularity (see methods and **Fig. 2c**) to detect single (monad) and double (dyad) spores, as the former are not easily distinguished from unsporulated cells by microscopy. We found 88% of wild yeast to sporulate in complete absence of nutrients. Only 0.6% of the domesticated isolates did so, and when sporulating they produced fewer asci (**Fig. 2d**). Neither wild nor domesticated yeasts capable of sporulating produced complete tetrads in water, forming only monads and dyads. This is consistent with spore numbers being controlled by carbon source availability^44^. To validate that domestication indeed abolished or impaired sporulation, we tracked sporulation in water across the never domesticated sister-species *S. paradoxus* (*n*=12 strains from 4 clades) and found it to be similar to that of wild *S. cerevisiae* (*n*=23) and much superior to that of domesticated *S. cerevisiae* (*n*=49) (**Supplementary fig. 2c**).

The capacity of wild yeast strains to sporulate and survive as spores in complete absence of external nutrients is important, as extreme starvation undoubtedly is a common challenge in natural habitats. We conclude that natural selection has maintained a remarkable capacity of wild yeasts to enter and survive in a haploid spore state, while domestication abolished or impaired this core organismal property. Because the domesticated clades have exchanged little or no genetic material, this abolishment of sporulation must have occurred independently multiple times during the yeast history.

### Domestication impaired sporulation through aneuploidies and sporulation gene loss

Because sporulation variation was overwhelmingly due to additive genetic effects (**Fig. 3a**, narrow sense heritability, *h*^2^=0.89 and 0.88, respectively at 24 and 72 h), we next probed the underlying genetic architecture and first focused on macroscopic genomic changes. Genome homozygosity has been reported to promote meiosis and sporulation^45^, but disregarding the wild yeasts, which are all completely homozygous, we found no correlation to sporulation (**Fig. 3b**). In fact, some populations (Sake, French Guiana, Wine/European *S. boulardii* subclade) had lost their sporulation capacity, despite high homozygosity. Because domesticated strains often carried ploidy variations, we next probed whether poly- or aneuploidy impaired sporulation. Tetraploid and triploid yeasts sporulated as often as diploid, and produced only marginally fewer asci at later time point (**Fig. 3c**). In contrast, aneuploidy accounted for some of the early and late sporulation variation (**Fig. 3c**). Multiple aneuploidies were worse than a single aneuploidy (**Fig. 3c**), and chromosome gain were as bad as chromosome loss. We propose that the presence of extensive chromosomal aneuploidies explains the poor sporulation of the domesticated Ale and African beer clades, as sporulation genes in these strains appeared to be free of loss-of-function nucleotide variants (see below).

**Figure 3.**
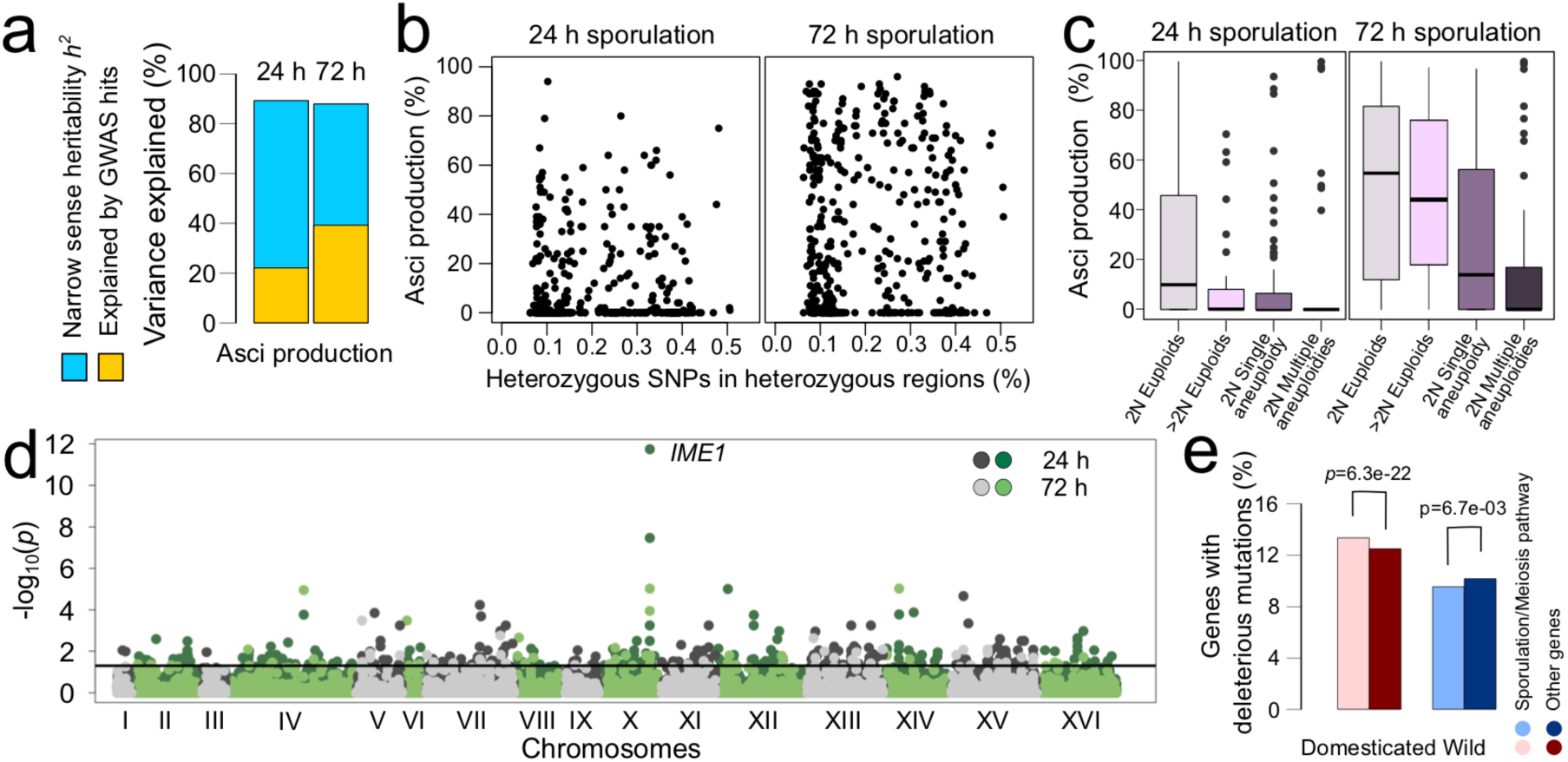
Domestication impaired sporulation through aneuploidy and loss of sporulation genes. **a**, Sporulation variance explained by genetics (narrow sense heritability) and additive effects of detected GWA hits. **b**, Sporulation efficiency after 3 days in standard sporulation environment (KAc) as a function of heterozygosity, estimated as heterozygote fraction of heterozygotic regions, after 24 h (*left panel*) and 72 h (*right panel*). **c**, *Left panel*: Comparing euploid diploids with polyploids (Mann-Whitney: *p*=2.9e-3), single and multiple aneuploidies (Mann-Whitney: *p*=1.7e-13, euploid vs. aneuploid; *p*=2.6e-2, single vs. multi aneuploidies) at 24 h. *Right panel*: Comparing euploid diploids with polyploids (Mann-Whitney: *p*>0.05), single and multiple aneuploidies (Mann-Whitney: *p*=6.3e-14, euploid vs. aneuploid; *p*=1.5e-2, single vs. multi aneuploidies) at 72 h. Box: IQR. Whiskers: 1.5x IQR. **d**, Manhattan plot showing the position of the GWAS hit across the genome with the sporulation master regulator *IME1* indicated. Black horizontal line indicates the threshold for significance (FDR α=0.05). **e**, Fraction of sporulation and non-sporulation genes carrying predicted loss-of-function mutations in domesticated and wild strains. Fisher’s exact test: *p*=1.7e-5 (domesticated vs. wild), *p*=6.3e-22 (sporulation vs. non-sporulation in domesticated) and 6.7e-3 (sporulation vs. non-sporulation in wild).

We next probed species-wide sporulation effects from common (MAF>5%) SNPs, indels and gains and losses of genes by GWAS and identified 384 sporulation variation markers (**Fig. 3d**). These were enriched (χ2, *p*=3.9e-05) in known sporulation genes (11%). The master regulator of meiosis, *IME1*, emerged as the primary sporulation determinant, with six previously unknown variants^46^ associating to this trait (**Supplementary fig. 3a**). Two promoter (A-325C, C-181T) and three coding (N311Y, E316V and E316G) new variants all associated with impaired sporulation (**Supplementary fig. 3b**), and predominantly occurred in domesticated lineages. A common (27%), derived H78R variant associated with enhanced sporulation in some domesticated yeasts (**Supplementary fig. 3c**).

Because sporulation effects of rare or clade specific variants evade GWAS detection^47^, we instead traced their effects by pinpointing likely loss-of-function variants in known sporulation and meiosis genes (*n*=251; **Supplementary table S3**) and through known sporulation QTNs^46,48,49^. Meiotic genes often carried loss-of-function variants in domesticated lineages, but rarely in wild yeasts (**Fig. 3e**). Abolished sporulation in the domesticated Sake and *S. boulardii* clades coincided with loss-of-function mutations in *RIM15* and *SPO74* (Sake), and in *MND1* and *SHE10* (*S. boulardii*) respectively. The negative effect of *RIM15* loss on sporulation has been experimentally validated^21^. In French Dairy and French Guyana clades sporulation loss was associated with loss-of-function variants in the key meiotic master regulators *NDT80* and *IME1*, as well as in the sporulation associated *OSW2, MND2, SPR6, DOM34, AQY1* (French Dairy) and *MPC54, DIT2, BMH2, DON1, NUD1* and *AQY1* (French Guiana). Nine domesticated Wine/European strains sporulated efficiently, but had premature stop codons (*SPO13* G86*; *SPO12* K103*) and early frameshifts in the key tetrad formation genes *SPO12* or *SPO13*^50^ (**Supplementary Fig. 4**) and only formed dyads.

Previously identified sporulation QTNs were mostly too rare in the sequenced global yeast population to affect sporulation variation overall, with e.g. *TAO3* (E1493Q), *SPO74* (C16A), *MKT1* (G30D) present in 2, 0 and 2 isolates respectively. Sporulation QTNs in *IME1* (L325M and A548G), *MCK1* (C1112A), *RSF1* (D181G), *PMS1* (A122G) and *FKH2* (A1498C) were more common (from 29 to 420 isolates with homozygous variants) but failed to associate with sporulation, consistent with their effects being restricted by epistasis to only a few genetic backgrounds^46^. Likely epistasis was e.g. found for *HOS4* (A1384G) and *SET3* (A1783T); while the 89 *SET3* (A1783T) homozygotes sporulated better overall (median: +130%), *HOS4* A1384G associated with enhanced sporulation only in *SET3* (A1783T) homozygotes.

### Quiescence survival is explained by population structure and local adaptation rather than domestication

An alternative life cycle strategy for yeasts encountering nutrient scarcity is to enter a quiescent G0 state^28^, which is also a complex trait that vary across genetic backgrounds^51^. We therefore probed whether domesticated isolates compensated for sporulation loss by evolving a superior quiescence life span. Encountering extreme starvation (water), wild yeasts all pursued the sporulation strategy, whereas domesticated and unclassified isolates (*n*=734) near uniformly entered quiescence. Quiescence survival (chronological life span; CLS) varied massively (0.5% to 96.8% survival after 20 days), but were on average much shorter than survival in the spore state. It is thus clearly an inferior survival strategy during nutrient scarcity. French Guiana and African Cocoa yeasts survived remarkably long in quiescence, while French dairy and African beer strains died rapidly. Aneu- and polyploidy both associated with an impaired quiescence life span under extreme starvation, similarly to what has been observed for the yeast replicative life span^52^. A single aneuploidy was as bad as multiple and chromosome gains were as bad as losses. Euploid polyploids died faster than diploids (36% vs. 62% median survival after 20 days; **Fig. 4a**).

**Figure 4.**
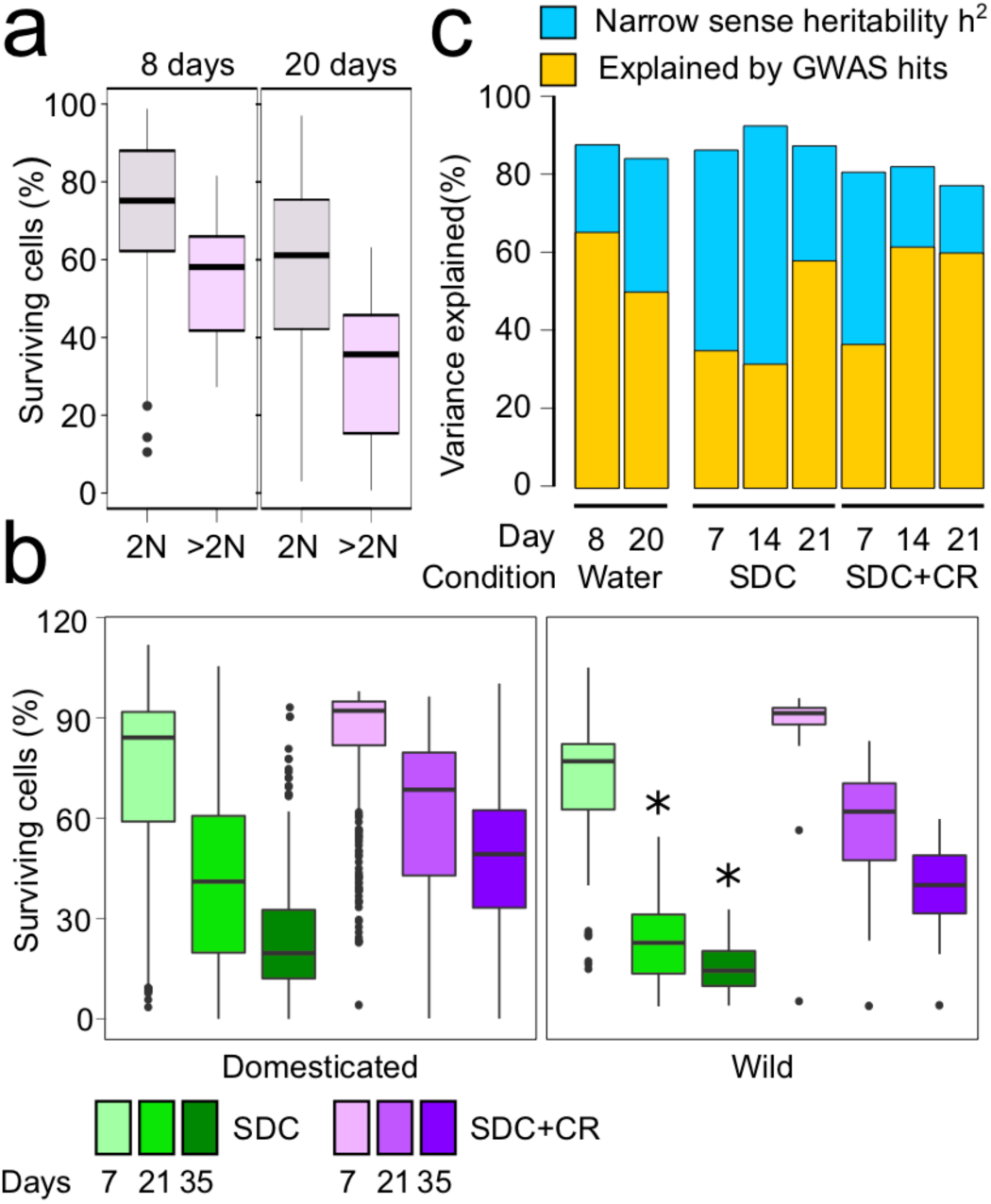
Domestication weakly extends yeast quiescence survival. **a**, Survival in water of euploid diploids and euploid polyploids 8 (upper panel) and 20 (lower panel) days after entry into quiescence. Mann-Whitney: *p*=6.7e-8 (8 days) and 3.2e-9 (20 days). Box: IQR. Whiskers: 1.5x IQR. **b**, Survival of wild and domesticated isolates at 7, 21 and 35 days after entry into quiescence following growth in rich synthetic complete medium (SDC, 2 % dextrose) or calorie restriction (SDC, 0.5 % dextrose). Significant Mann-Whitney (FDR, *α*=0.05) comparisons of wild vs. domesticated isolates are indicated by asterisks. Box: IQR. Whiskers: 1.5x IQR. **c**, Survival variance explained by genetics (broad sense heritability) and by additive effects of detected GWA hits.

We next compared the quiescence survival of wild and domesticated yeasts (*n*=572, all euploid diploid), in carbon exhausted synthetic media in which not even wild yeast sporulates. We allowed cells to reach quiescence by exhausting sugars either using rich synthetic complete medium (SDC) or a calorie restricted such medium. After 7, 21, and 35 days we found nearly equal quiescence survival for wild and domesticated yeasts, with a small but significant advantage for domesticated yeasts in the richest environment (SDC, at days 21 and 35) that best resemble domestication niches (**Fig. 4 panel B**). Rather than by domestication, survival variation under quiescence was explained by population stratification and therefore either by drift or local adaptation. Thus, wild Taiwanese and domesticated French Dairy yeasts died rapidly (<30% survivors after 7 days), while French Guiana, Mexican agave and West African Cocoa yeasts were the longest living (>60% survivors after 21 days) (**Supplementary fig. 4**). French Dairy carried loss-of-function mutations in *SIC1*, which is required for entry into quiescence^53^, whereas French Guiana yeasts had a defective allele of *PCL7*, which represses quiescence^54,55^ (see **Supplementary table S3** for the complete list of candidate genes).

Given the high trait heritability (**Fig. 4c**, *h*^2^ = from 0.77 to 0.90), we next probed alleles controlling survival under quiescence by GWAS. We identified 292 variants which were mostly (*n*=177) time and environment specific. Loss function mutations in the cell cycle regulator *WHI2* was associated to shortened quiescence lifespan in all environments. Furthermore, 14 common (MAF>16%) variants in the minor succinate dehydrogenase (*SDH1b*), assisting mitochondrial electron transfer, were associated to shortened quiescence life span under all but extreme starvation. The succinate dehydrogenase complex probably controls yeast viability through its involvement in the TCA cycle and in the mitochondrial respiratory chain, which must be operational to promote survival upon nutrient exhaustion^56–58^.

Calorie restriction extended the quiescence lifespan for all clades, except for in the extremely long lived West African Cocoa yeasts (**Supplementary Fig. 5**). An intergenic variant far (−2065) upstream *HPF1* and the common (*n*=69) presence of a highly diverged non-reference *HPF1* allele, both extended survival under calorie restriction at all quiescence time points and result in being the top GWAS hits. We have recently shown intragenic repeat expansions in *HPF1* itself to be a key regulator of yeast quiescence life span, through controlling buoyancy and thereby oxygen exposure and altering redox signalling^59^. This finding suggest that such mode of regulation may be common through yeast populations given that intragenic repeat numbers evolve rapidly^60,61^.

## Discussion

Domestication impairing yeast sporulation represents a truly seismic change to the biology of the organism. Firstly, loss of capacity to enter the yeast spore state, which offers long term survival when nutrients are scarce^30,31^, leaves domesticated cells instead dependent on entering the G0 quiescence stage. Entry into quiescence also slows ageing and intrinsic death^62^, but much less so than sporulation (**Fig. 2d**). Moreover, domesticated yeasts only marginally compensated for the sporulation loss by evolving extended quiescence life spans (**Fig. 4b**). Domestication therefore caused a massive net drop in capacity to survive nutrient scarcity. This is likely to be detrimental in wild microhabitats, where nutrient scarcity is the rule rather than the exception for yeasts that lack motility and are unable to relocate when local nutrients are consumed. Secondly, sporulation entails transitioning from a higher to a lower ploidy state. Domesticated yeasts will have no or a much reduced capacity to achieve this shift. Because a lower ploidy state confer faster asexual reproduction in many environments^63^, this is likely to translate into a fitness cost. Third, loss of sporulation means loss of the capacity to form gametes and meiotic recombination. Recombination also occurs under yeast mitosis but at vastly lower rates^64,65^. Thus, domestication near abolished or impaired the yeast capacity to combine beneficial variants into one genotype, to cleanse otherwise healthy genomes from deleterious mutations^66,67^, to prevent their catastrophic accumulation through a ratchet mechanism^68^ and to generate a sufficiently broad diversity of genotypes to fully exploit and adapt to multi-faceted wild niches^69^. Sexual recombination accelerates yeast adaptation^70^ and the loss of sexual recombination is likely to be particularly costly in constantly changing wild niches where fast adaptation is key. Rare subclades of domesticated clades, in particular of the Wine/European lineage, are known to sometimes escape from their domestication niches and to survive as feral isolates in the wild. This is likely thanks to these strains retaining, or having re-acquired, a life cycle closer to that of wild yeast. Indeed, sporulation of feral subclades in the Wine/European clade is notably more efficient than that of other sub-clades (**Supplementary fig. 2b**). Whereas purifying selection retains a functional sporulation machinery in all wild isolates, there is little doubt that non-feral domesticated strains with their impaired sporulation capacity, are strongly selected against in the wild.

Because the yeast domestication syndrome is shared across genetically distinct, independently domesticated clades, it is unlikely to reflect the retention of ancestral trait that have been lost in wild lineages^71^. Instead, it likely represents convergent evolution where the convergence was driven by domestication. Such a convergence could have both adaptive and neutral explanations. First, the domestication syndrome re-oriented both carbon and nitrogen metabolism from better respiratory to better fermentative growth. This can easily be rationalized as adaptation. Domestication niches, such as grape must with a 20% glucose/fructose content, are often rich in easily fermented sugars. Ethanol production, which is the principle trait selected for in most domestication environments, depends on efficient fermentation of these sugars. Respiration of the sugar, or of the produced ethanol, is both counterproductive to high ethanol content and discouraged by high sugar and, later, low oxygen. The better beer yeast fermentation of malt sugars to ethanol^23^, is widely held to reflect domestication driven adaptation. In contrast, concentrated sugar is relative rare in wild niches and competition for it is fierce, with concentrations unlikely to often exceed the 0.5% threshold at which sugar fermentation kicks in. In contrast to in large industrial tanks where the oxygen is rapidly depleted, oxygen is also often freely available in natural yeast niches, allowing respiration. Second, we found sporulation genes to associate to clade specific loss-of-function variants in genes private to sporulation and meiosis. This is consistent with a neutral drift model, where asexual proliferation in domestication niches relaxes selection on superfluous sporulation and meiosis genes, leaving them free to accumulate mutations. Many of these variants affected the key meiosis specific transcription factors *IME1* and *NDT80* that are absolutely required for sporulation^72^, underscoring that there is little selective pressure to retain sporulation intact in domesticated isolates. Sporulation requires respiration^73,74^. Relaxed selection on sporulation may therefore indirectly lower the selection also for respiration, and this could partially account for the worse respiratory growth in domesticated lineages. Niche specific loss of superfluous genes, leading to pseudogenization and ultimately genome reduction, has been suggested to be the predominant driver of natural yeast evolution^21,75,76^. Domestication may simply be seen as directing and accelerating this pseudogenization process. A notable direct side-effect of relaxed selection on the sexual life cycle is that correct pairing and segregation of chromosomes during meiosis becomes irrelevant. This reduces selection against chromosomal rearrangements, aneuploidies and odd polyploidies that disturb these processes, allowing their accumulation in domesticated strains^20,77^. This creates a knock-on effect, as their accumulation in domesticated strains reinforces the loss of the sexual cycle. Besides domestication, two events stand out in *S. cerevisiae* evolution since the *S. paradoxus* split. First, the out-of-China dispersal separated East Asian wild isolates from other yeast clades, driving the earliest and most profound genetic differentiations^20,78,79^. Second, hybridization of some *S. cerevisiae* strains to wild *S. paradoxus*, followed by repeated backcrossing to the *S. cerevisiae* parental lineage, later created *S. cerevisiae* genomes, such as the Alpechin clade, with massive (up to 5%) amounts of introgressed *S. paradoxus* DNA^20^. Both events affect asexual growth. However, we found neither to have altered the yeast life cycle, the core organism-level property of the species. Domestication therefore stands out as the most dramatic event in yeast evolution, being unique in having reprogrammed a central aspect of its organismal biology. Fundamental consequences of domestication on life cycles are not unique to yeast: domestication made banana (*Musa sp*.) seedless and parthenocarpic^80^ and abolished seed and flower production in garlic (*Allium sativum*)^81^. In contrast to these species, however, yeast is not only industrially important but serves as one of our most widely used model species. Because our understanding of yeast biology is near entirely based on studies of domesticated, or predominantly domesticated, strains^24^, this calls for reflection. How much of what we know about this key model species reflects, domestication rather than natural biology and how much has this influenced our understanding of biology at large?

## Material and methods

### Yeast strains

We used 987 globally sampled *S. cerevisiae* isolates, thereby capturing the spectrum of geographic and ecological sources, including both wild and domestication niches^20^. Strains were recently sequenced, their single nucleotide variant based population structure established, and most isolates assigned to one of 26 phylogenetic clades. Because individual domesticated isolates can escape into wild niches and individual wild isolates invade domestication niches, we assigned isolates as wild (*n*=58) or domesticated (*n*=574) based on the origin dominating (>66%) in their clade. Strains were labelled “unassigned” (*n*=355) if neither criteria were reached; however, we note that unassigned strains behave as domesticated for most traits here studied. For *asexual growth*, estimates all *n*=987 isolates were retained. For estimates of *quiescence survival in plain water, n*=485 domesticated, *n*=52 wild and *n*=289 unassigned isolates with sufficient cell counts, no aggregating and no sporulating isolates were retained. For estimates of *quiescence survival in other environments, n*=342 domesticated, *n*=48 wild and *n*=182 unassigned euploid diploid isolates with sufficient cell counts, no aggregation and not sporulating were retained. For *sporulation* estimates *n*=503 domesticated, *n*=54 wild and *n*=293 unassigned non-haploid isolates capable of growth in the pre-sporulation medium were retained. Subsets of isolates were used in follow-up sporulation experiments, as indicated in text.

### Asexual growth

Clonal populations of all isolate were pre-cultivated as colonies in a 1536 array on top of solid agar (2%) plates containing synthetic complete (SC) medium (0.14% YNB, 0.5% ammonium sulfate, 0.077% Complete Supplement Mixture (CSM, Formedium), 2% (w/v) glucose and pH buffered to 5.8 with 1% (w/v) succinic acid and 0.6% (w/v) NaOH) for 72 h at 30°C. For alternative nitrogen sources, CSM was excluded and ammonium sulfate limited to 30 mg N/L to prevent nitrogen storage and subsequent growth on stored nitrogen. Pre-cultures were subsampled with short pins and subsamples transferred as 1536 arrays to experimental plates containing the medium of interest (background medium as above). For alternative carbon sources or concentrations, 2% glucose was replaced by the indicated carbon source at the indicated concentration. For alternative nitrogen sources, ammonium sulfate and CSM were replaced by the alternative nitrogen source at growth yield limiting concentrations (30 mg N/L). Cultivation using the Scan-o-matic platform^82^ was performed in thermostatic cabinets at 30°C and constant, high humidity. Cells in each colony were counted at 20 min intervals using high-quality desktop scanners and Scan-o-matic 2.2 (https://github.com/Scan-o-Matic/scanomatic), with exact pixel intensity calibration and transformation into cell counts. Population size growth curves were established, smoothed and transformed into population doubling times (*d*) and total cell yields (*y*), as described^82^. To account for spatial variation within and between plates, fixed genotypes (the diploid hybrid DBVPG6756xYPS128) were included in every 4th of the 1536 colony positions on each plate, and their log_2_ *d* and *y* measures were used to normalize the corresponding *d* and *y* measures of adjacent colonies. We estimated the mean normalized log_2_ *d* and *y* in each environment for domesticated and wild isolates and called significant differences using a Mann-Whitney test with an FDR-adjusted *α*=0.05.

### Sporulation efficiency and spore viability

We measured sporulation both in traditional sporulation medium and plain, isotonic water. *Sporulation in traditional sporulation medium:* Isolates were pre-cultivated in yeast peptone dextrose (YPD; 2% dextrose, 1% yeast extract, 2% peptone, 2% agar) before being diluted 1:50 into 10 mL of pre-sporulation media (YPA; 2% potassium acetate, 1% yeast extract, 2% peptone) and grown 48 hours at 30°C (shaking: 250 rpm). Pre-sporulated cells were transferred to sporulation media (2% KAc) into 250 ml flasks at a 5:1 volume/medium ratio and a final optical density (OD_600_) of 0.6. Flasks were kept at 23 °C and shaken at 250 rpm. To estimate sporulation efficiency, >100 cells/sample were counted at 24 and 72 h post-transfer to sporulation medium using an optical microscope (Zeiss Axio Lab.A1). Sporulation efficiency was estimated as the number of asci divided by the total cell count. To estimate survival in the spore state, 300 µL of sporulated cells were sampled at the indicated time points, washed 2x with water and resuspended in 100 µL of dissection buffer (sorbitol 2 M, EDTA 10 mM, sodium phosphate 100 mM, zymolase 1 mg/mL) for 1 h at 37 °C. 32 tetrads per isolate and time point were dissected on YPD plates with the micro-dissecting microscope SporePlay+ (Singer Ltd, UK). Plates were incubated 3 days at 30°C and surviving spores was estimated as the fraction of dissected spores capable of expanding asexually to form visible colonies.

#### Sporulation in plain water

Isogenic cultures were grown overnight in liquid YPD in 96 well plates, diluted 1:100 in 200 µL of fresh YPD in a 96 well plate and incubated for 24 hours at 30°C. 20 µL of each culture was transferred to a new 96 well plate containing 180 µL of phosphate saline buffer, representing isotonic water. Cells were washed (2x) twice with PBS to remove remaining YPD, and finally resuspended in 200 µL of PBS (day 0). For each sample and time point, we counted 10,000 cells in a FACS Calibur flow cytometer (Becton Dickinson) using the forward (FSC; cell size) and side (SSC; cell granularity) scatter to distinguish two cell populations: unsporulated, quiescent cells and spores present as smaller and less granular monads. We estimated the sporulation efficiency as the number of asci formed divided by the total cell count. Sporulation among detected asci was confirmed in a wild strain (AKN) by quantification of the promoter activity of *SPS2*, induced late in sporulation, using a *SPS2*-GFP fusion protein reporter^83^. Finally, the sporulation efficiency was visually confirmed in a subset of samples by microscopy (as above).

### Survival in quiescence

We estimated quiescence survival in plain water and in calorie rich and calorie restricted synthetic defined medium in two distinct sets of experiments. *Quiescence survival in plain water:* Isogenic cultures from *n*=485 domesticated, *n*=52 wild and *n*=289 unassigned isolates were independently grown overnight in liquid YPD in 96 well plates, diluted 1:100 in 200 µL of fresh YPD and incubated for 24 hours at 30°C. 20 µL of each culture were then transferred to 96 well plates containing 180 µL of phosphate saline buffer (PBS). Cells were washed (2x) and resuspended in 200 µL of PBS (day 0) for the rest of the experiment. Plates were sealed with adhesive aluminium foil and kept at 30°C. *Quiescence survival in SDC and calorie restricted SDC:* Isogenic cultures of *n*=342 domesticated, *n*=48 wild and *n*=182 unassigned euploid diploid isolates were independently grown overnight in liquid SDC (2% dextrose, 0.675% yeast nitrogen base (Formedium), 0.088% complete amino acid supplement (Formedium), pH buffered to 6.0 with 2.5 M NaOH), diluted 1:100 in 200 µL of either fresh SDC or calorie restricted SDC (0.5% dextrose instead of 2%) in 96 well plates. Plates were incubated at 30°C and the cells were kept in saturated media for the whole experiment. Aging was considered to start at saturation of the cultures, 72 h post-inoculation^84^.

#### Quiescence survival measurements (all environments)

chronological life span was measured as viable cells (%) by flow cytometry based on the uptake of the fluorescent molecules propidium iodide (PI) and YO-PRO-1 iodide (YP). Propidium iodide and YO-PRO-1 are membrane-impermeable nucleic acid binding molecules that enter into necrotic but not into alive cells. Therefore, non-fluorescent cells are alive, while fluorescent cells are not. YO-PRO-1 penetrates also into apoptotic cells^85,86^. At each indicated aging time point, 5 µL of cells were transferred into 100 µL of staining solution (Phosphate Buffer Saline + 3 µM propidium iodide (Sigma-Aldrich) + 200 nM YP (Invitrogen)) in a 96 well plate and incubated for 5 min in the dark at 30°C. Samples were analysed on a FACS-Calibur flow cytometer (Becton Dickinson) using a High Throughput Sampler (Becton Dickinson) device to process 96 well plates and detect fluorescence with FL-1 (YP; 488/10 nm excitation, 530/30 nm emission) and FL-3 (PI; 488/10 nm excitation, 670 nm emission) channels. Experiments were run in a single replicate and data were normalized to 4 internal controls distributed in each plate. 10,000 cells per sample were analysed.

### Candidate gene lists

The candidate gene list for sporulation variants was constructed by including all genes belonging to seven GO:TERM categories: GO:0000710 (meiotic mismatch repair), GO:0007126 (meiotic nuclear division), GO:0007127 (meiosis I), GO:0030435 (sporulation resulting in formation of a cellular spore), GO:0030437 (ascospore formation), GO:0051321 (meiotic cell cycle) and GO:0060903 (positive regulation of meiosis I). The candidate gene list for chronological life span was constructed as in Matecic *at al.* 2010^87^.

### Loss-of-function variants

We used SnpEff^88^ (v.4.1) and SIFT^89^ (Sorting Intolerant from Tolerant; v.5.2.2) to predict single base variant effects. An in-house script was used to estimate small frameshifting indels of multiples other than three. Likely losses of gene function were called when such frameshifts, premature stop codons, or loss of either the predicted start or stop codon were found. Assuming loss-of-function variants to be completely dominated by functional variants, we called gene loss-of-function only when such variants were homozygotic.

### Genome wide association

The VCF file available for the sequenced strains^20^ was converted and filtered using plink (v1.9) with MAF>0.05 for loss-of-function SNPs, gene presence/absence variants and copy number variants (called as single versus multiple copies). Non biallelic SNPs were retained but for these only the most frequent of the minor alleles was used. Loss-of-functions, gene presence/absence and copy number variants were based on the non-reduntant pangenome described in Peter et al. 2018^20^ and manually added to the plink SNPs files at simulated genomic positions. GWA, heritability and the genome inflation factor were all calculated estimated using the R function `fast-lmm`. GWA correction for population structure was performed for SNPs with a minor allele frequency, MAF>0.5%. To correct for population structure when estimating heritability and the genome inflation factor, genetic variants used as markers were included.

### Statistics

Unless otherwise stated, statistical 2-group comparisons were performed in R.3.1.1, using the function `wilcox.test,` a 2-sided Mann-Whitney test and FDR correction for multiple hypothesis testing at *α*=0.05. Effect sizes were calculated using the R function `cliff.delta` which reduces the influence of outliers.

## Acknowledgements

The authors thank Folkert van Werven, Gilles Fisher and Bertrand Llorente for the helpful discussions and Barak Cohen for the kind gift of the *SPS2::GFP* tagged strains. This work was supported by Agence Nationale de la Recherche (ANR-11-LABX-0028-01, ANR-13-BSV6-0006-01, ANR-15-IDEX-01, ANR-16-CE12-0019 and ANR-18-CE12-0004) and the Swedish Research Council (2014-6547, 2014-4605, 2018-03638 and 2018-03453).

## Supplementary materials

**Table S1. Differences in stress resilience.** Significant (1-sided Mann-Whitney test; FDR α=0.05) differences in resilience to stresses between wild and domesticated or wine feral and other wine isolates.

**Table S2. Differences in nutrient usage.** Significant (1-sided Mann-Whitney test; FDR α=0.05) differences in doubling time (*d*) and yield (*y*) between wild and domesticated isolates. Effect sizes are included.

**Table S3. Lists of candidate genes.** We obtained lists of candidate genes (see Methods) by combining GO:TERM categories for sporulation and meiosis (sporulation list) and from Matecic *at al.* 2010^87^ (quiescence genes) respectively.

**Supplementary data S1**

All measured phenotypes, sporulation GWAS hits, chronological life span GWAS hits, yield GWAS hits, doubling time GWAS hits.

**Supplementary data S2**

Variant matrix for GWAS generated by Plink in .bed, .bim and .fam formats. The matrix contains all biallelic positions known for the sequenced isolates with MAF > 5% as well as LOF (encoded as “1” for predicted LOF and “2” for predicted functional gene), presence and absence (encoded as “1” for gene presence and “2” for gene absence gene) and CNVs (encoded as “0”, “1”, “2” respectively for absence, 0.5–1 copy and multiple copies).

---

**Supplementary Figure 1.**
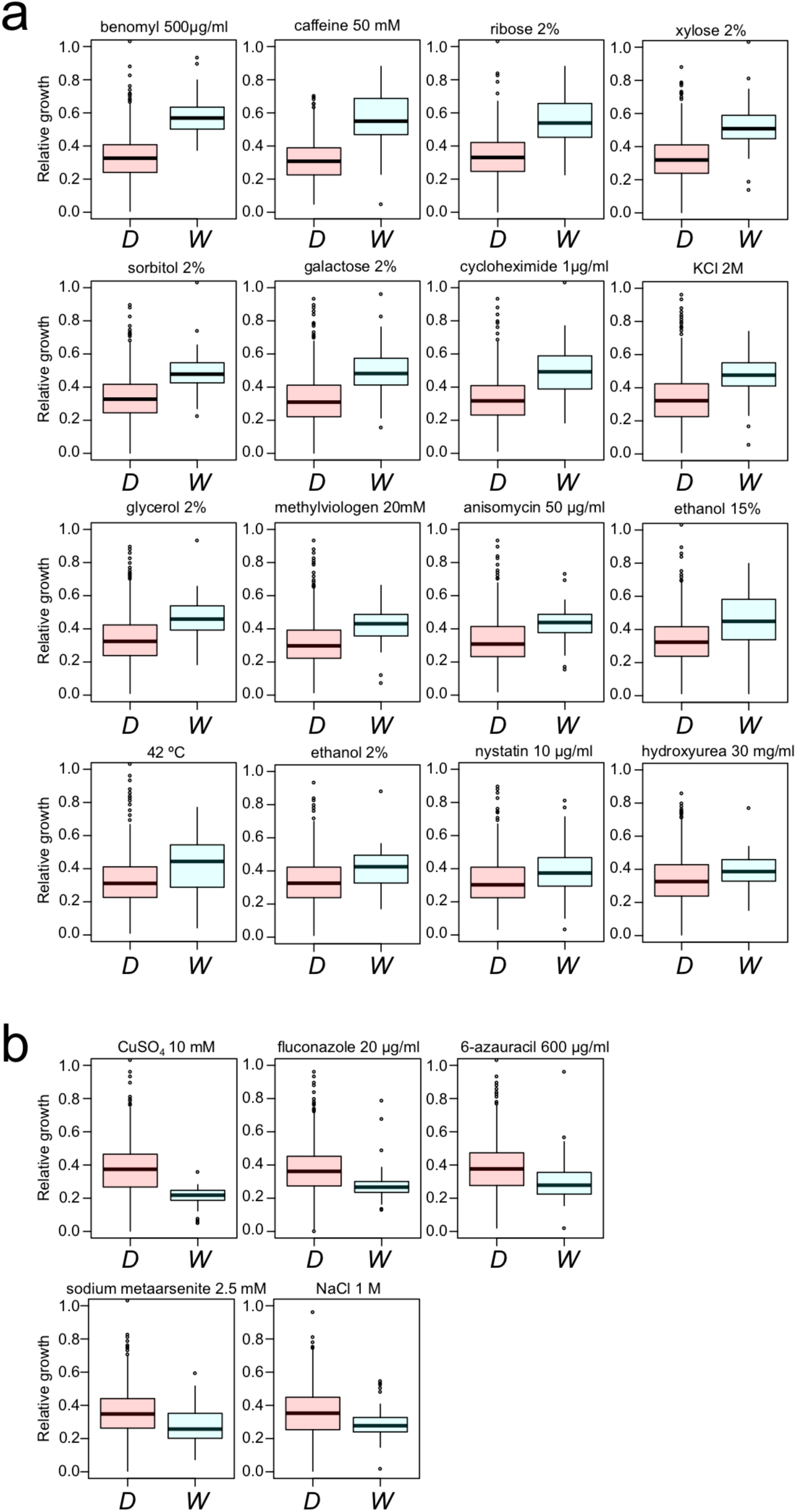
Wild clades grow better in stress. Growth data were taken from Peter *et al*. 2018^20^. **a**, Boxplot of growth in the 16 environments in which wild (*W*) are superior to domesticated strains (*D*). **b**, Boxplot of growth in the five environments in which domesticated are superior to wild strains. Box: IQR. Whiskers: 1.5x IQR.

**Supplementary Figure 2.**
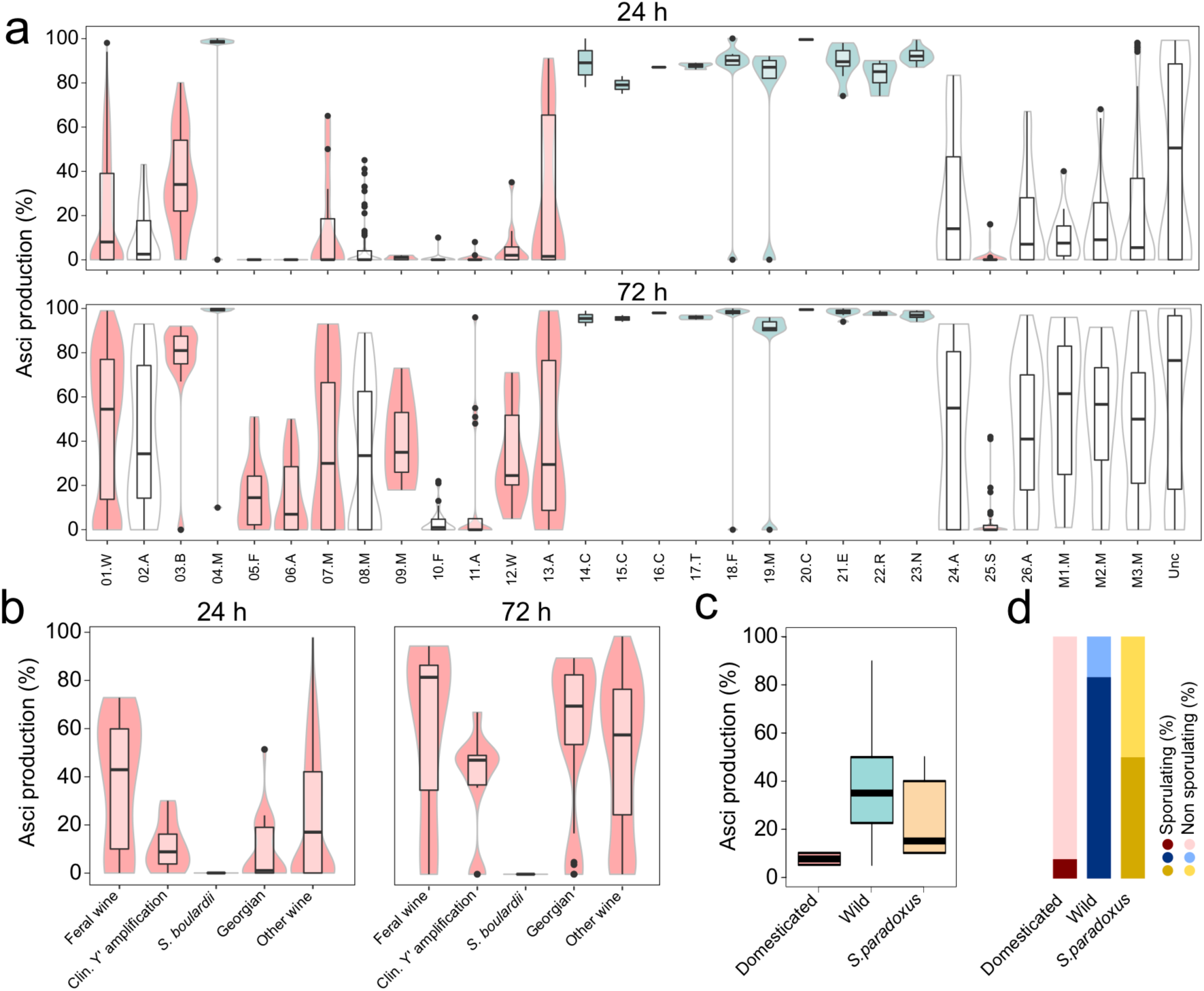
Asci production varies across domesticated but not wild yeast clades. **a**, Distribution of the asci production after 24 h (upper panel) and 72 h (lower panel) in standard experimental sporulation environment (KAc) for each phylogenetic clade. Colours indicate domesticated (red) or wild (blue), unassigned clade are left white. Box: IQR. Whiskers: 1.5x IQR. **b**, Distribution of the asci production at 24 h and 72 h for the four subclades within the Wine/European clade. Feral isolates regained a high sporulation efficiency. The *S. boulardii* subgroup has completely lost the ability to sporulate. Box: IQR. Whiskers: 1.5 X IQR. **c**, Sporulation ability in water. *S. cerevisiae* and *S. paradoxus* wild strains sporulate well, but *S. cerevisiae* domesticated isolates do not. Boxplot of the distribution of asci production (only sporulating isolates, i.e. asci production > 0) after 8 days in water. Box: IQR. Whiskers: 1.5x IQR. **d**, Fraction of *S. cerevisiae* domesticated, wild and *S. paradoxus* isolates able to sporulate (i.e. asci production > 0 after 8 days) in water.

**Supplementary Figure 3.**
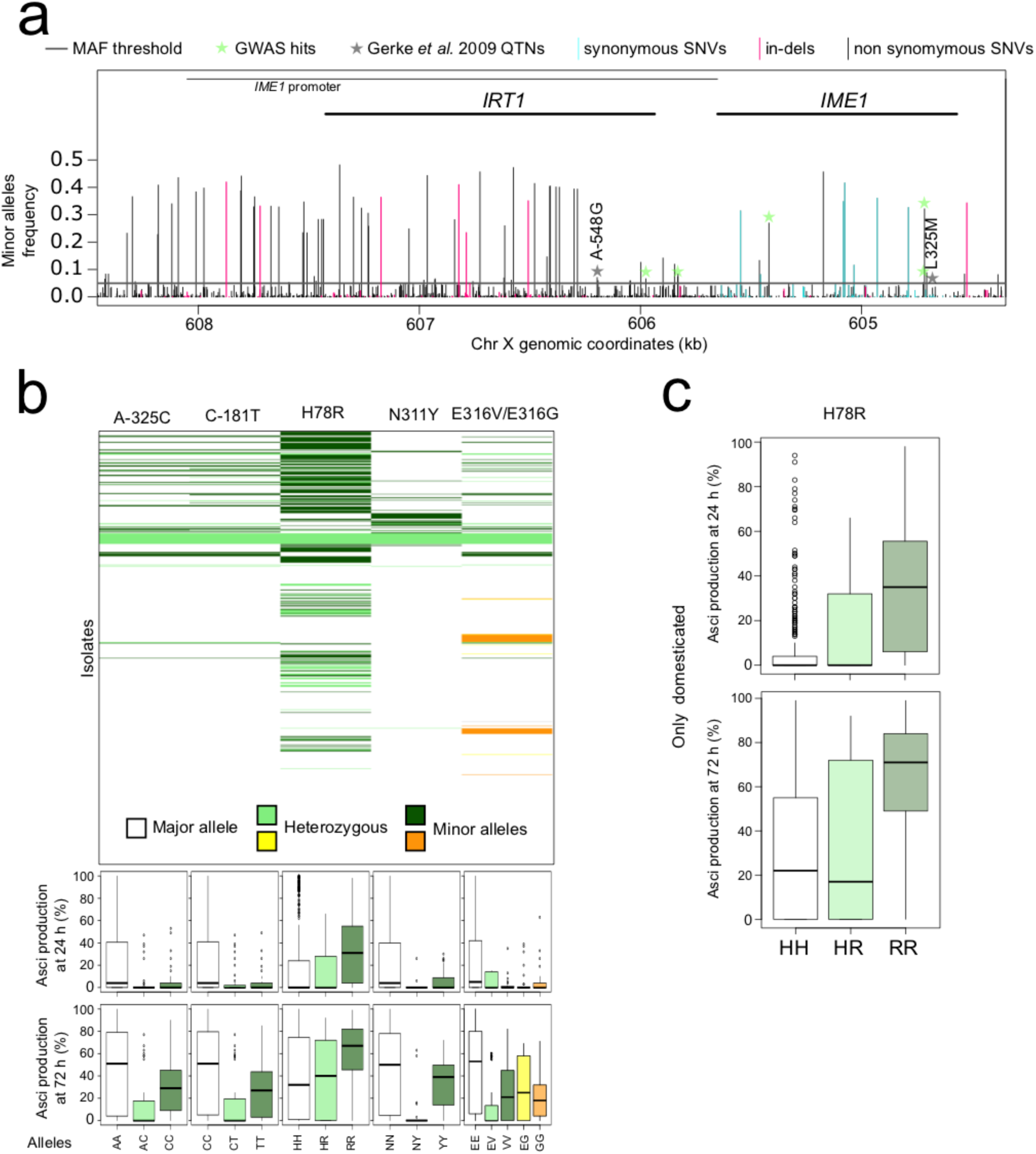
*IME1* variants control yeast sporulation variation. **a**, Schematic representation of the meiotic regulator *IME1* locus and the upstream *IRT1* gene, which overlaps the *IME1* promoter. GWAS revealed five SNPs (green stars) in the *IME1* and its promoter associate to yeast sporulation after 24 h in standard KAc. None of these correspond to the SNPs identified in Gerke *et al* 2009^46^ (grey stars), which do not pass the MAF>0.05. **b**, *Upper panel:* Distribution of *IME1* GWAS sporulation hits across the sequenced strain collection (*y*-axis, ordered as in the tree phylogeny reported in Peter *et al* 2018^20^). Several SNPs are linked. *Lower panel:* Sporulation of strains homo- or heterozygotic for the five *IME1* SNPs associated to sporulation. Box: IQR. Whiskers: 1.5x IQR. All minor SNPs except H78R associate to poor spore production. **c**, H78R is restricted to a subset of domesticated clades and promotes spore production. Box: IQR. Whiskers: 1.5x IQR.

**Supplementary Figure 4.**
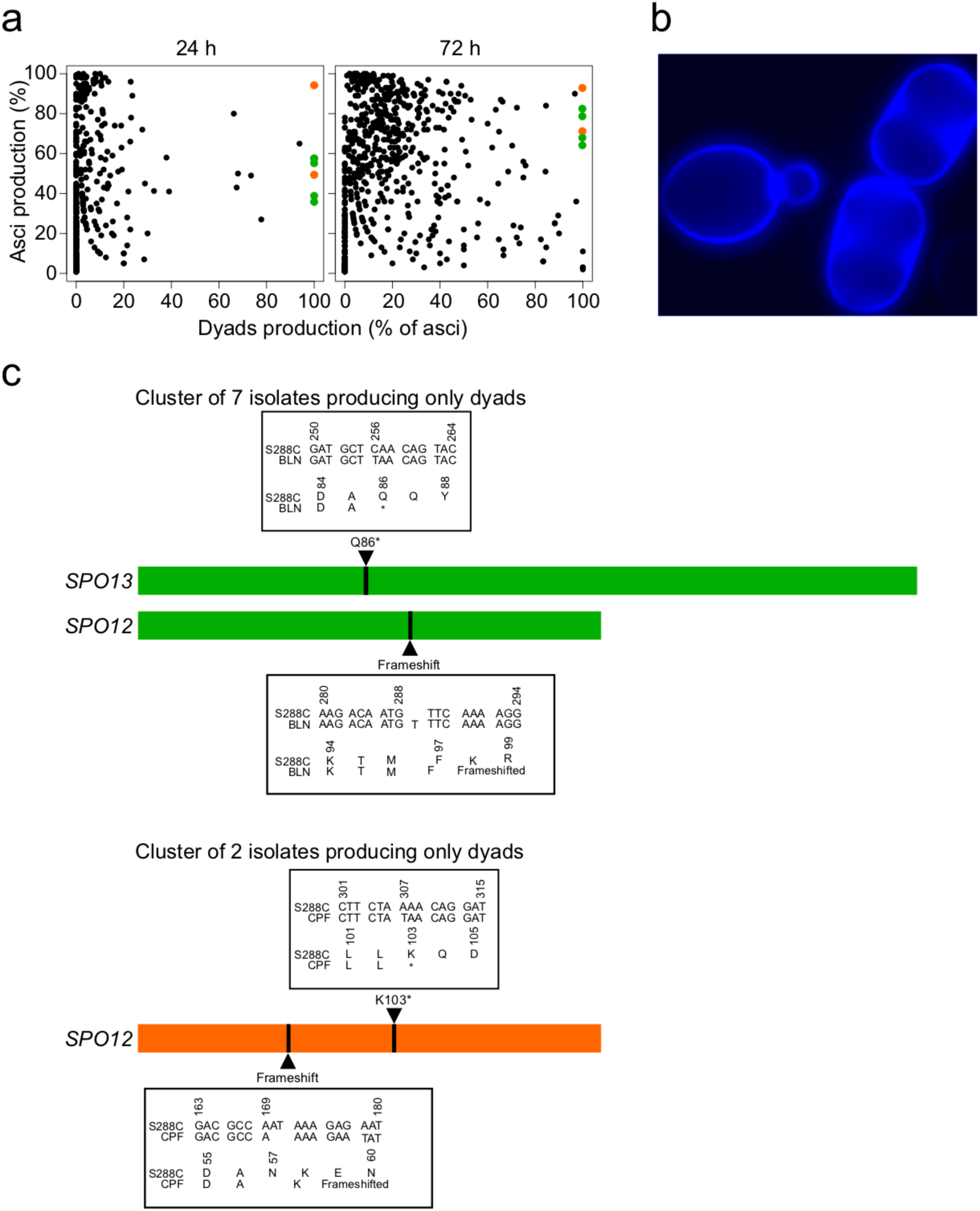
Loss-of-function variants in *SPO1*2 and *SPO13* trigger dyads-only spore production. **a**, Sporulation (asci production, *y*-axis) after 3 days in standard experimental sporulation environment (KAc) and the percentage of produced asci that are dyads (*x*-axis), after 24 h (*left panel*) and 72 h (*right panel*). Nine strains, belonging to two clusters (indicated in green and orange) efficiently enter sporulation but interrupt meiosis cycle after meiosis I, only producing dyads. **b**, Micrograph of asci produced in strain BNL, after 72 h of sporulation, with staining of the spore wall (blue). Only dyads are produced. **c**, Schematic representation of amino acid sequences for *SPO12* and *SPO13* showing position of homozygous loss-of-function variants that trigger dyads-only production during sporulation.

**Supplementary Figure 5.**
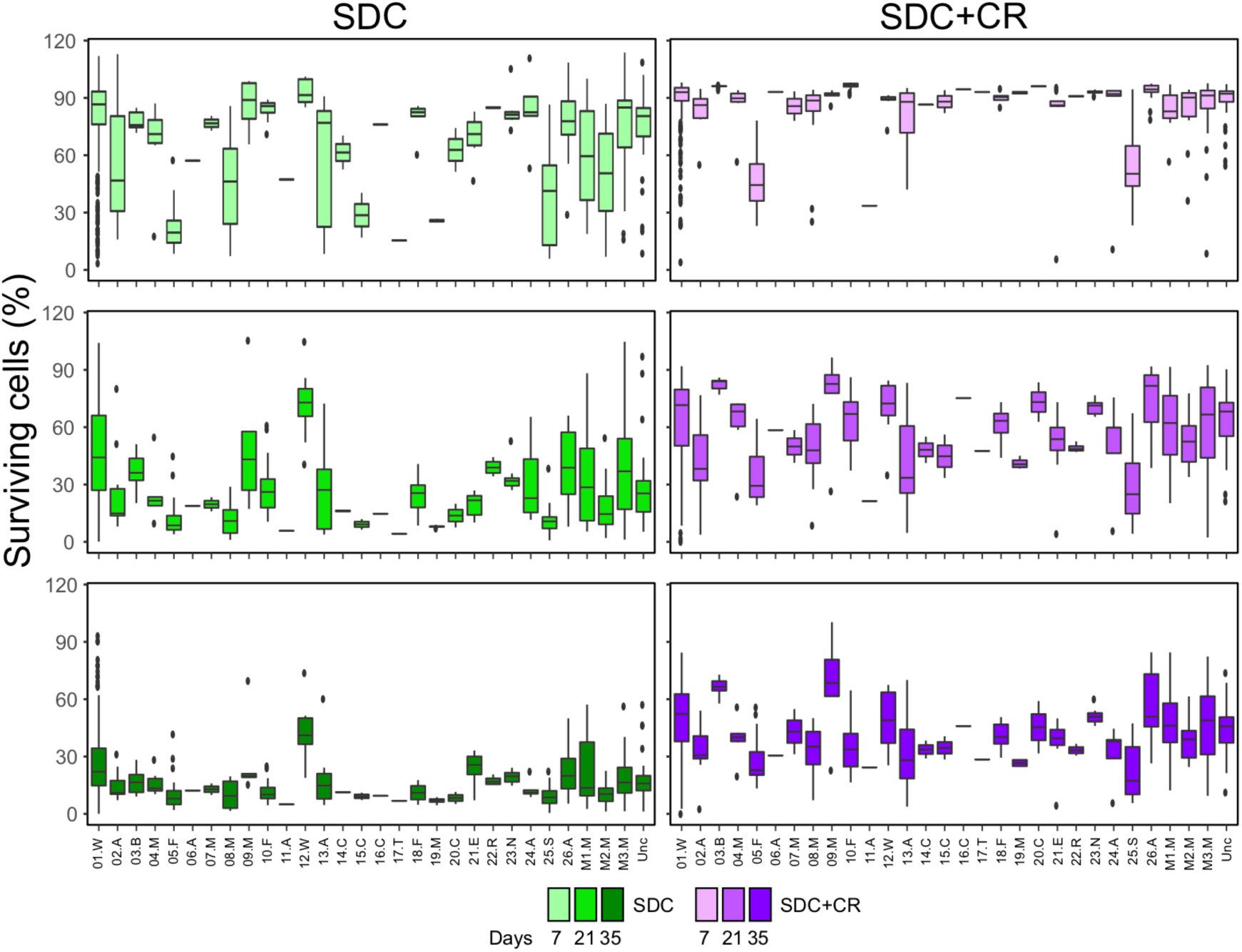
Population stratification determines yeast quiescence survival. Boxplots showing surviving cells (%) for each clade in each quiescence condition and time point. French Dairy (5.F) yeast has the lowest survival while French Guiana (10.F), Mexican agave (09.M) and West African Cocoa (12.W) have the highest. Box: IQR. Whiskers: 1.5 X IQR.

